# Aedes anphevirus (AeAV): an insect-specific virus distributed worldwide in *Aedes aegypti* mosquitoes that has complex interplays with *Wolbachia* and dengue virus infection in cells

**DOI:** 10.1101/335067

**Authors:** Rhys Parry, Sassan Asgari

## Abstract

Insect specific viruses (ISVs) of the yellow fever mosquito *Aedes aegypti* have been demonstrated to modulate transmission of arboviruses such as dengue virus (DENV) and West Nile virus by the mosquito. The diversity and composition of the virome of *Ae. aegypti*, however, remains poorly understood. In this study, we characterised Aedes anphevirus (AeAV), a negative-sense RNA virus from the order *Mononegavirales*. AeAV identified from *Aedes* cell lines were infectious to both *Ae. aegypti* and *Aedes albopictus* cells, but not to three mammalian cell lines. To understand the incidence and genetic diversity of AeAV, we assembled 17 coding-complete and two partial genomes of AeAV from available RNA-Seq data. AeAV appears to transmit vertically and be present in laboratory colonies, wild-caught mosquitoes and cell lines worldwide. Phylogenetic analysis of AeAV strains indicates that as the *Ae. aegypti* mosquito has expanded into the Americas and Asia-Pacific, AeAV has evolved into monophyletic African, American and Asia-Pacific lineages. The endosymbiotic bacterium *Wolbachia pipientis* restricts positive-sense RNA viruses in *Ae. aegypti*. Re-analysis of a small RNA library of *Ae. aegypti* cells co-infected with AeAV and *Wolbachia* produces an abundant RNAi response consistent with persistent virus replication. We found *Wolbachia* enhances replication of AeAV when compared to a tetracycline cleared cell line, and AeAV modestly reduces DENV replication *in vitro*. The results from our study improve understanding of the diversity and evolution of the virome of *Ae. aegypti* and adds to previous evidence that shows *Wolbachia* does not restrict a range of negative strand RNA viruses.

## Importance

The mosquito *Aedes aegypti* transmits a number of arthropod-borne viruses (arboviruses) such as dengue virus and Zika virus. Mosquitoes also harbour insect-specific viruses that may affect replication of pathogenic arboviruses in their body. Currently, however, there are only a handful of insect-specific viruses described from *Ae. aegypti* in the literature. Here, we characterise a novel negative strand virus, Aedes anphevirus (AeAV). Meta-analysis of *Ae. aegypti* samples showed that it is present in *Ae. aegypti* mosquitoes worldwide and is vertically transmitted. *Wolbachia* transinfected mosquitoes are currently being used in biocontrol as they effectively block transmission of several positive sense RNA viruses in mosquitoes. Our results demonstrate that *Wolbachia* enhances the replication of AeAV and modestly reduces dengue virus replication in a cell line model. This study expands our understanding of the virome in *Ae. aegypti* as well as providing insight into the complexity of the *Wolbachia* virus restriction phenotype.

## Introduction

The yellow fever mosquito *Aedes aegypti* is a vector of medically important viruses with worldwide distribution within the tropical and subtropical zones (1). *Ae. aegypti* is the principal vector of both dengue virus (DENV) and Zika virus (ZIKV) (Family: *Flaviviridae*) with estimates suggesting up to 390 million incidences of DENV infections a year (2), and approximately 400,000 cases of ZIKV during the 2015–2016 Latin American ZIKV outbreak (3).

The ability of mosquitoes to transmit viruses is determined by a complex suite of genetic and extrinsic host factors (4–6). One developing area is the contribution of insect-specific viruses (ISVs), demonstrated not to replicate in mammalian cells, in the vector competence of individual mosquitoes (7, 8). ISVs can suppress the anti-viral RNAi response as shown in Culex-Y virus (CYV) of the *Birnaviridae* family (9), or enhance the transcription of host factors; cell fusing agent virus (CFAV) (Family: *Flaviviridae*) infection of *Ae. aegypti* Aa20 cells upregulates the V-ATPase-associated factor RNASEK allowing more favourable replication of DENV (10). ISVs have also been shown to suppress or exclude replication of arboviruses; prior infection of *Aedes albopictus* C6/36 cells and *Ae. aegypti* mosquitoes with Palm Creek virus (PCV) (Family: *Flaviviridae*) has been shown to supress replication of the zoonotic West Nile virus (WNV) and Murray Valley encephalitis virus (Family: *Flaviviridae*) (11, 12). Also, it has recently been demonstrated in *Aedes* cell lines that dual infection with Phasi Charoen-like virus (Family: *Bunyaviridae*) and CFAV restricts the cells permissivity to both DENV and ZIKV infection (13).

Metagenomic and bio-surveillance strategies have proved invaluable in describing the virome diversity of wild-caught *Culicinae* mosquitoes (14, 15). To date, six ISVs have been identified and characterised from wild-caught and laboratory *Ae. aegypti;* CFAV (16, 17), Phasi Charoen-like virus (Family: *Bunyaviridae*) (18), Dezidougou virus from the Negevirus taxon (19), Aedes densoviruses (Family: *Parvoviridae*) (20) and the unclassified Humaita-Tubiacanga virus (HTV) (21). Recently, transcriptomic analysis of wild-caught *Ae. aegypti* mosquitoes from Bangkok, Thailand and Cairns, Australia suggested possible infection of the mosquitoes with up to 27 insect-specific viruses, the majority of which currently uncharacterized (22). This represents a narrow understanding of the diversity of the circulating virome harboured by *Ae. aegypti* mosquitoes.

In this study, we identified and characterised a novel negative-sense RNA *Anphevirus*, putatively named Aedes anphevirus (AeAV), from the order *Mononegavirales* in *Ae. aegypti* mosquitoes. According to the most recent International Committee on Taxonomy of Viruses (ICTV) report (23), Xīnchéng mosquito virus (XcMV), assembled as part of a metagenomic analysis of *Anopheles sinensis* mosquitoes in Xīnchéng China, is the only member of the *Anphevirus* genus and closely related to members of *Bornaviridae* and *Nyamiviridae* (24). Originally thought to only encode for four ORFs, presence of a number of closely related viruses to XcMV from West African *Anopheles gambiae* mosquitoes (15) and West Australian *Culex* mosquitoes (25) suggests that members of this taxon encode for six ORFs with a genome size of approximately 12kb.

The endosymbiotic bacterium *Wolbachia pipientis* was first shown to restrict RNA viruses in *Drosophila melanogaster* (26, 27). Transinfection of *Wolbachia* into *Ae. aegypti* restricted DENV and Chikungunya virus (Family: *Togaviridae*) replication in the host (28). In *Ae. aegypti* Aag2 cells, stably transinfected with a proliferative strain of *Wolbachia* (wMelPop-CLA), the endosymbiont restricts CFAV (29), but has no effect on the negative sense Phasi Charoen-like virus (Family: *Bunyaviridae*) (30). In addition to characterising AeAV, we also studied the effect of *Wolbachia* on AeAV replication and co-infection of AeAV and DENV in *Ae. aegypti* cells.

## Results

### Identification and assembly of the full Aedes anphevirus (AeAV) genome from *Wolbachia*-infected *Aedes* cells

During replication of RNA viruses in *Ae. aegypti* mosquitoes, the RNA interference (RNAi) pathway cleaves viral dsRNA intermediates into 21nt short interfering RNAs (vsiRNAs) (31, 32). Using this fraction of reads from RNA-Seq data, it is possible to *de novo* assemble virus genomes (21, 33).

The previously sequenced small RNA fraction of embryonic *Ae. aegypti* Aag2 cells and Aag2 cells stably infected with *Wolbachia* (wMelPop-CLA strain) (34) was trimmed of adapters, filtered for 21nt reads and *de novo* assembled using CLC Genomics Workbench with a minimum contig length of 100nt. The resulting contigs were then queried using BLASTX against a local virus protein database downloaded from the National Centre for Biotechnology Information (NCBI). In the Aag2.wMelPop-CLA assembly, four contigs between 396-1162nt were found to have amino acid similarity (E value 9.46E-51) to proteins from two closely related *Mononegavirales* viruses: Culex mononega-like virus 1 (CMLV-1) and Xīnchéng mosquito virus (XcMV); the type species for the *Anphevirus* genus. No contigs from the *Wolbachia* negative *Ae. aegypti* Aag2 dataset showed any similarity to CMLV-1 or XnMV.

Subsequent RT-PCR analysis between RNA samples from Aag2 and Aag2. wMelPop-CLA cell lines indicated that this tentative virus was exclusive to the Aag2.wMelPop-CLA cell line (Fig. 1A). We hypothesised that the presence of any putative virus may have been the result of contamination during *Wolbachia* transinfection. The *Wolbachia* wMelPop-CLA strain was isolated from the *Ae. albopictus* cell line RML-12 and transinfected into Aag2 (35) and the *Ae. albopictus* C6/36 (C6/36. wMelPop-CLA) cell lines (36). RT-PCR analysis of RNA extracted from RML-12 and C6/36 cells, as well as the *Ae. aegypti* cell line Aa20, showed that the putative virus was present only in RML-12 cells (Fig. 1A).

**Figure 1.**
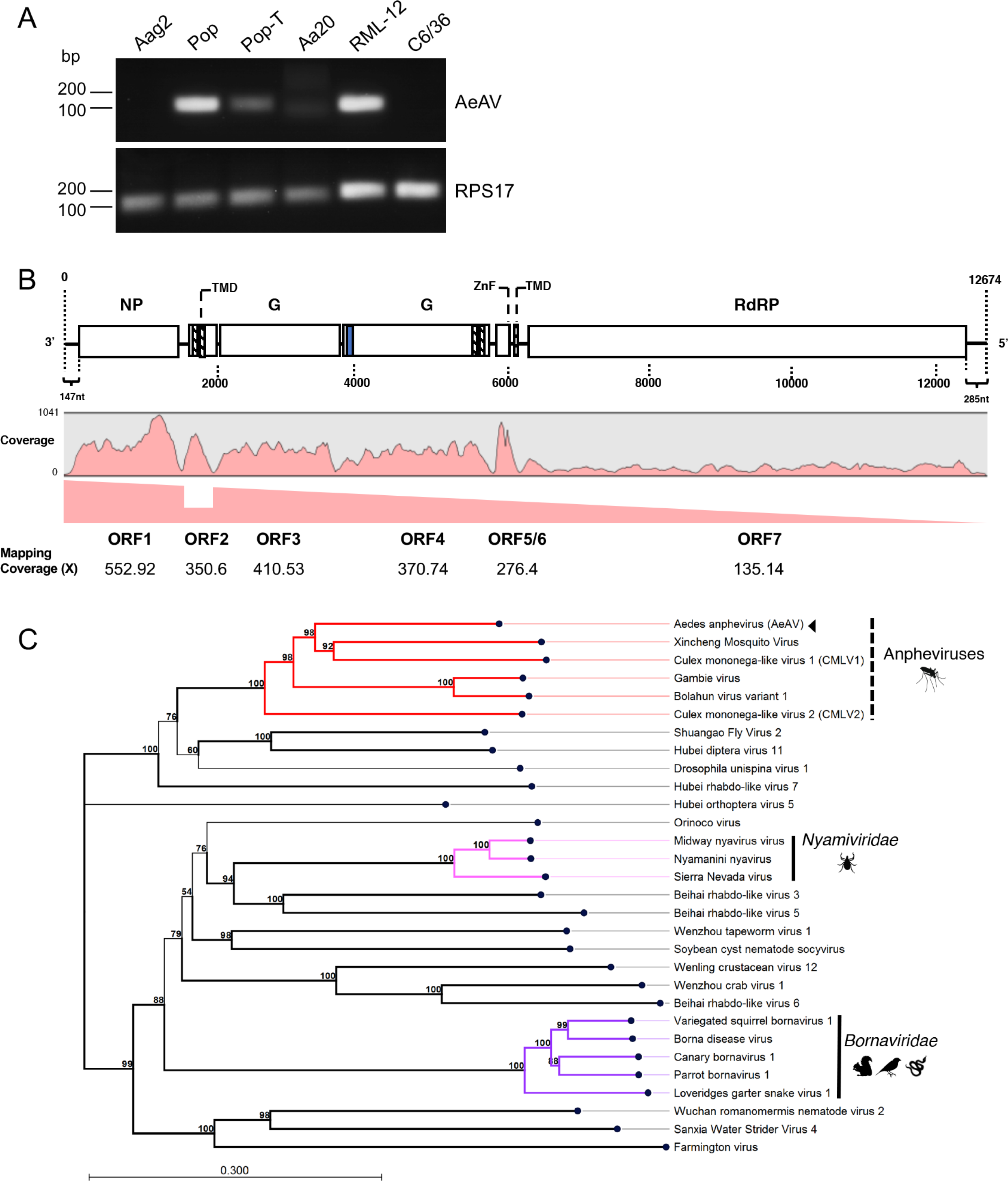
Presence of Aedes anphevirus (AeAV) in insect cell lines, and genome organisation and phylogeny of the virus. **A)** RT-PCR analysis of *Aedes* cell lines Aag2, Aag2. *w*MelPop-CLA (Pop), *w*MelPop-CLA.Tet (Pop-T), Aa20, RML-12, and C6/36 for the presence of AeAV. RPS17 was used as a loading control. **B)** Genome organisation of the Cali, Colombia AeAV genome strain and subgenomic gene transcription profile. Transmembrane domains (TMD) are depicted as boxes with dashed lines and signal peptide is depicted as a blue box. NP, nucleoprotein; G, glycoprotein; ZnF, zinc-like finger; RdRP, RNA dependent RNA polymerase. **C)** AeAV is a member of the *Anphevirus* genus (red), related to members of the *Nyamiviridae* (pink) and *Bornaviridae* (purple) in an unassigned family within the order *Mononegavirales*. A multiple sequence alignment of the RNA dependent RNA polymerase and the mRNA capping domain was used to create a Maximum likelihood phylogeny. The phylogeny is arbitrarily rooted. 1000 bootstraps were performed and branches with bootstrap values greater than 85% are highlighted. Branch length represent expected numbers of substitutions per amino acid site. Genbank protein accession numbers are Bolahun virus variant 1 (AOR51366.1), Culex mononega-like virus 1 (CMLV1) (ASA47369.1), Culex mononega-like virus 2 (CMLV2) (ASA47322.1), Gambie virus (AOR51379.1), Xincheng Mosquito Virus (XcMV) (YP_009302387.1), Borna disease virus (YP_009269418.1), Canary bornavirus 1 (YP_009268910.1), Loveridges garter snake virus 1 (YP_009055063.1), Parrot bornavirus 1 (AEG78314.1), Variegated squirrel bornavirus 1 (SBT82903.1), Midway nyavirus (YP_002905331.1), Nyamanini nyavirus (YP_002905337.1), Sierra Nevada virus (YP_009044201.1), Soybean cyst nematode socyvirus (YP_009052467.1), Farmington virus (YP_009091823.1), Beihai rhabdo-like virus 3 (APG78650.1), Beihai rhabdo-like virus 5 (YP_009333422.1), Beihai rhabdo-like virus 6 (YP_009333413.1), Drosophila unispina virus 1 (AMK09260.1), Hubei diptera virus 11 (YP_009337182.1), Hubei orthoptera virus 5 (YP_009336728.1), Hubei rhabdo-like virus 7 (YP_009337121.1), Orinoco virus (ANQ45640.1), Sanxia Water Strider Virus 4 (YP_009288955.1), Shuangao Fly Virus 2 (AJG39135.1), Wenling crustacean virus 12 (YP_009336618.1), Wenzhou Crab Virus 1 (YP_009304558.1), Wenzhou tapeworm virus 1 (YP_009342311.1), Wuchan romanomermis nematode virus 2 (YP_009342285.1).

To recover the remainder of the virus genome, transcriptome RNA-Seq data from RML-12.wMelPop-CLA and C6/36.wMelPop-CLA cells were downloaded (37, 38) and *de novo* assembled with automatic bubble and word sizes using CLC Genomics Workbench. BLASTN analysis of assembled contigs indicated that a complete 12,940 contig from the C6/36. *w*MelPop-CLA cells and two contigs (9624 and 3487nt) from RML-12. *w*MelPop-CLA were almost identical (99-100% pairwise nucleotide identity) to the virus like contigs assembled from Aag2. *w*MelPop-CLA. We were then able to use this reference to recover the full genomes from Aag2. *w*MelPop-CLA and RML-12. *w*MelPop-CLA consensus mapping to this reference. To re-validate that AeAV was only present in Aag2. *w*MelPop-CLA cells, reads from the *Wolbachia*-negative Aag2 cells were mapped to the representative genome, and only four reads were identified in the data. The result from here and the RT-PCR analysis above (Fig. 1A) also confirm that the virus found in the *Wolbachia* transinfected cells originate from RML-12 cells in which *w*MelPop-CLA was originally transinfected and subsequently transferred to other cell lines.

### Characterisation of Aedes anphevirus (AeAV)

AeAV genomes assembled in this study were between 12,455 to 13,011 nucleotides in length with a %GC content of 46.8% and encode for 7 non-overlapping ORFs (Fig. 1B). Phylogenetic analysis of the RNA-dependent RNA polymerase protein places AeAV within a well-supported clade of the unassigned *Anphevirus* genus, which are from the order *Mononegavirales* and closely related to members of *Bornaviridae* and *Nyamiviridae* (Fig. 1C).

All members of *Mononegavirales* have a negative-stranded RNA genome encapsidated within the capsid and the RNA polymerase complex (39). The RNA genome is used as the template by the RNA polymerase complex to sequentially transcribe discrete mRNAs from subgenomic genes. mRNA from each gene is capped and polyadenylated. To analyse the transcriptional activity of AeAV, we used the poly-A enriched RNA-Seq libraries prepared from the Cali, Colombia laboratory strain (40). Read mapping and coverage analysis of the AeAV genome showed that AeAV follows the trend of reduced transcriptional activity seen in other *Mononegavirales* species (41) with approximately 50% reduction between ORF1 and ORF2 but an increased transcription between ORF2 and ORF3 (Fig. 1B). The reduction in transcriptional activity of AeAV genes is conserved for each sequential ORF with the least transcriptional activity for ORF7/L protein, that is conserved in all AeAV strains in Poly(A) enriched RNA-Seq libraries (Fig. S1).

ORF1 of AeAV encodes a predicted 49kDa nucleoprotein with no transmembrane domains and closest pairwise amino acid identity (26%) to the nucleoprotein gene from Culex mononega-like virus 1 (CMLV-1) from *Culex* mosquitoes in Western Australia (25). Protein homology analysis using HHPred showed that ORF1 was a likely homolog of the p40 nucleoprotein of the Borna disease virus (Probability 98.66%, E-value: 7.1e-10). ORF2 encodes an 11kDa protein with two transmembrane domains in the N-terminus of the protein with no similarity to any proteins within the non-redundant protein database or homologs as predicted by HHPred. ORF3 and ORF4 encode putative glycoproteins, 64kDa and 72kDa, respectively. ORF3 has no pairwise amino acid similarity to any virus protein or homologs as per HHPred analysis. ORF4 was predicted to have a signal peptide in the N-terminus followed by a heavily O- and N-linked glycosylated outside region as well as two transmembrane domains in the C-terminus of the protein. ORF4 is most closely related to the glycoprotein from the Gambie virus identified from West African *An. gambiae* mosquitoes with 45% pairwise amino acid identity (15). Protein homology analysis predicted ORF4 to be a homolog of the Human Herpesvirus 1 Envelope Glycoprotein B (Probability 99.88%, E-value 2.2e-22).

The presence of a Zinc-like finger (ZnF) domain in a small ORF proximal to the L protein previously reported in closely related viruses (15) (Fig. 2A and B), was identified in AeAV based on sequence alignment (Fig. 1B). Re-analysis of putative ORFs from CMLV-1 and CMLV-2 (25) showed the presence of this GATA-like ZnF domain in both of these viruses and the genus type species XcMV identified from *An. sinesis* (24) (Fig. 2C).

**Figure 2.**
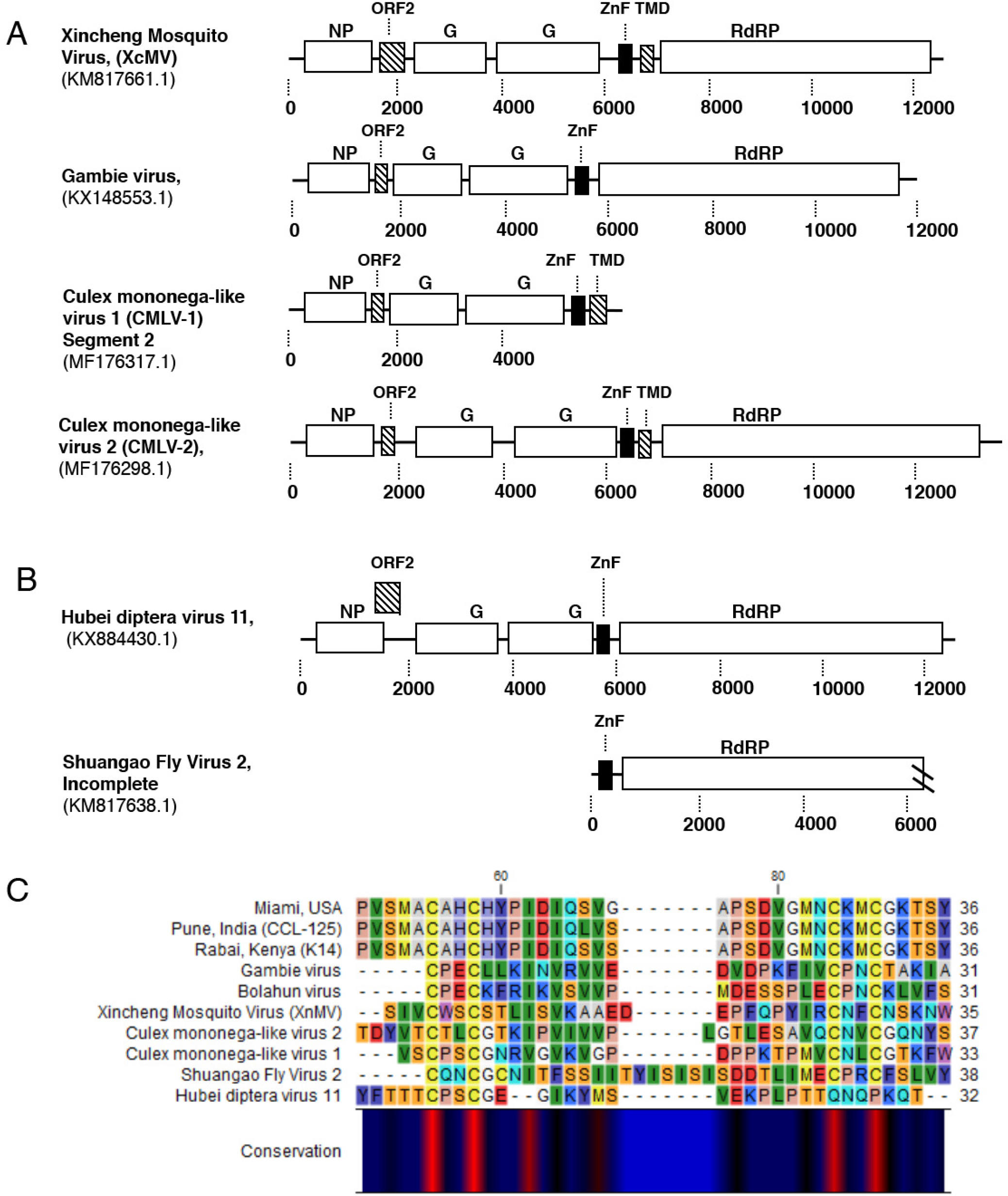
Conservation of GATA-like Zinc finger (ZnF) domain and small transmembrane domain containing protein between tentative members of the *Anphevirus* taxon. **A)** Genome orientation of previously discovered viruses within the *Anphevirus* taxon, and **B)** two viruses within a closely related clade. Predicted ORF encoding the ZnF domain is indicated by a black square. Predicted ORFs containing transmembrane domains are indicated by dashed lines. Genbank accession numbers are shown below virus name. NP, nucleoprotein; G, glycoprotein; ZnF, zinclike finger; RdRP, RNA dependent RNA polymerase. **C)** Alignment of predicted GATA-like ZnF protein sequence (C-X(2)-C-X(17-20)-C-X(2)-C) between three representative strains of AeAV (Miami, USA; Pune, India; Rabai, Kenya) and predicted ZnF domain proteins from Figure 2A and B.

ORF6 encodes for a small 4kDa protein that has a single transmembrane domain in the C-terminus. This protein was almost missed in the prediction of ORFs due to having only 37 amino acids, however, it has a strong transcriptional coverage in Poly-A datasets and exists in all assembled strains (Fig. 1B and Fig. S1). It was predicted to share no structural homology or amino acid identity with any previously reported peptide. In addition to this, we were able to identify small transmembrane domain containing proteins proximal to or overlapping with the ZnF protein in CMLV-1, CMLV-2 and XcMV (Fig. 2A), suggesting that this protein may be a conserved feature of anpheviruses.

ORF7 encodes for the 226kDa L protein, has 41% pairwise amino acid identity with the RNA dependent RNA polymerase from CMLV-1. Protein domain analysis of the L protein showed the highly conserved *Mononegavirales* RNA dependent RNA polymerase, mRNA capping domain and a mRNA (guanine-7-) methyltransferase (G-7-MTase) domain conserved in all L proteins in *Mononegavirales* (42).

### AeAV *cis*-regulatory elements

For identification of *cis*-regulatory elements in the AeAV genome, we used MEME (Multiple Em for Motif Elicitation) to search for overrepresented 5-50nt motifs (43). Using a 0-order Markov model, one 32nt motif 3’-UUVCUHWUAAAAAACCCGCYAGUUASAAAUCA-5’ was considered statistically significant (E-value: 4.2e-010). Importantly the motif was proximal to each predicted virus gene ORF, suggesting it may be a potential promoter (Fig. 3A and C). No motif was found between ORF 5 and 6 in AeAV suggesting that these two genes may be under the control of a single *cis*-regulatory element. Interestingly, the complement of this motif appeared twice on the anti-genome suggesting that it may be used in an anti-genome virus intermediate. We noticed that these motifs localised to partial palindromic repeats and predicted that they may form stable secondary RNA structures. Using RNAfold, we were able to visualise and predict the MFE structure 20nt upstream and downstream of the motif (44). All predicted *cis*-regulatory motifs formed partial or complete stable secondary stem loops and hairpins with high base-pair probabilities (Fig. 3B). The exception was predicted element 3, which is proximal to the second ORF; as this second gene is transcribed less than ORF3 irrespective of its similarity to the motif, the lack of a stable stem loop structure may be a novel transcriptional regulatory mechanism. The presence of two conserved homopolymeric triplets in the overrepresented motif is very similar to “slippery” −1 ribosome frame shifting (RFS) sites XXX YYY Z (X=A, G, U; Y=A, U; Z=A, C, U) (45). It has been previously demonstrated that similar ‘slippery’ sequence motifs followed by a predicted stem-loop structure is a feature of rhabdovirus gene overlap regions (46). In AeAV, this feature appears in the intergenic space and is unlikely to represent ribosomal frame shifting event and subsequent extension of a protein. We also searched for additional slippery motifs in the AeAV genome. The genomic context for each predicted “slippery motif” did not extend or produce additional ORFs.

**Figure 3.**
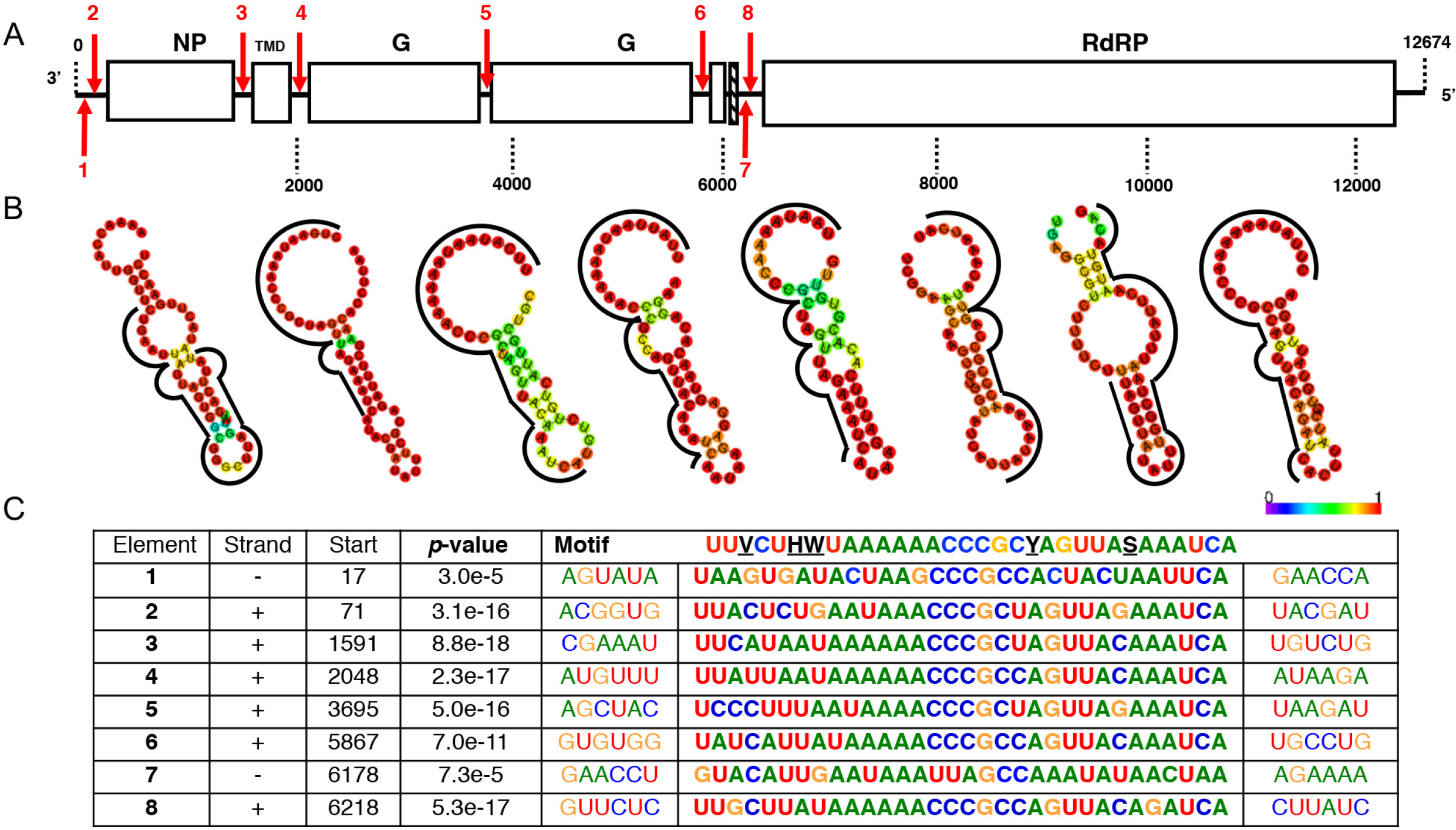
Aedes anphevirus (AeAV) *cis*-regulatory elements. **A)** Location and orientation of predicted *cis*-regulatory element in AeAV indicated by numbered red arrows; downwards indicating genome and upwards arrow indicating anti-genome. **B)** Predicted minimum free energy (MFE) RNA structure of the region surrounding the motif for each element using the RNAfold web server. Colour indicates probability of base-pairing and motif is indicated by the black line. **C)** Sequence of the conserved motif as predicted by MEME as well as location and the statistical confidence of the motif. Sequences are written 3’ to 5’ and anti-genome motif sequences 1 and 7 are depicted as reverse complement for visual clarity.

### AeAV infection is widespread in *Ae. aegypti* laboratory colonies, wild-caught mosquitoes and cell lines

Taking advantage of the currently published RNA-Seq data, we performed a meta-analysis of global incidence and genetic diversity of this virus. We were able to show that AeAV is ubiquitous in laboratory colonies, cell lines and wild-caught *Ae. aegypti* mosquitoes. During preparation of this manuscript a partial AeAV genome of 5313nt (Accession: MG012486.1) was deposited into NCBI nucleotide database from a study characterising the evolution of piRNA pathways across arthropods (47). Using our AeAV genome reference, we were able to complete the CDS portion of the genome and also 16 additional strains of AeAV with two additional incomplete genomes (Fig. 4) (Table S1).

**Figure 4.**
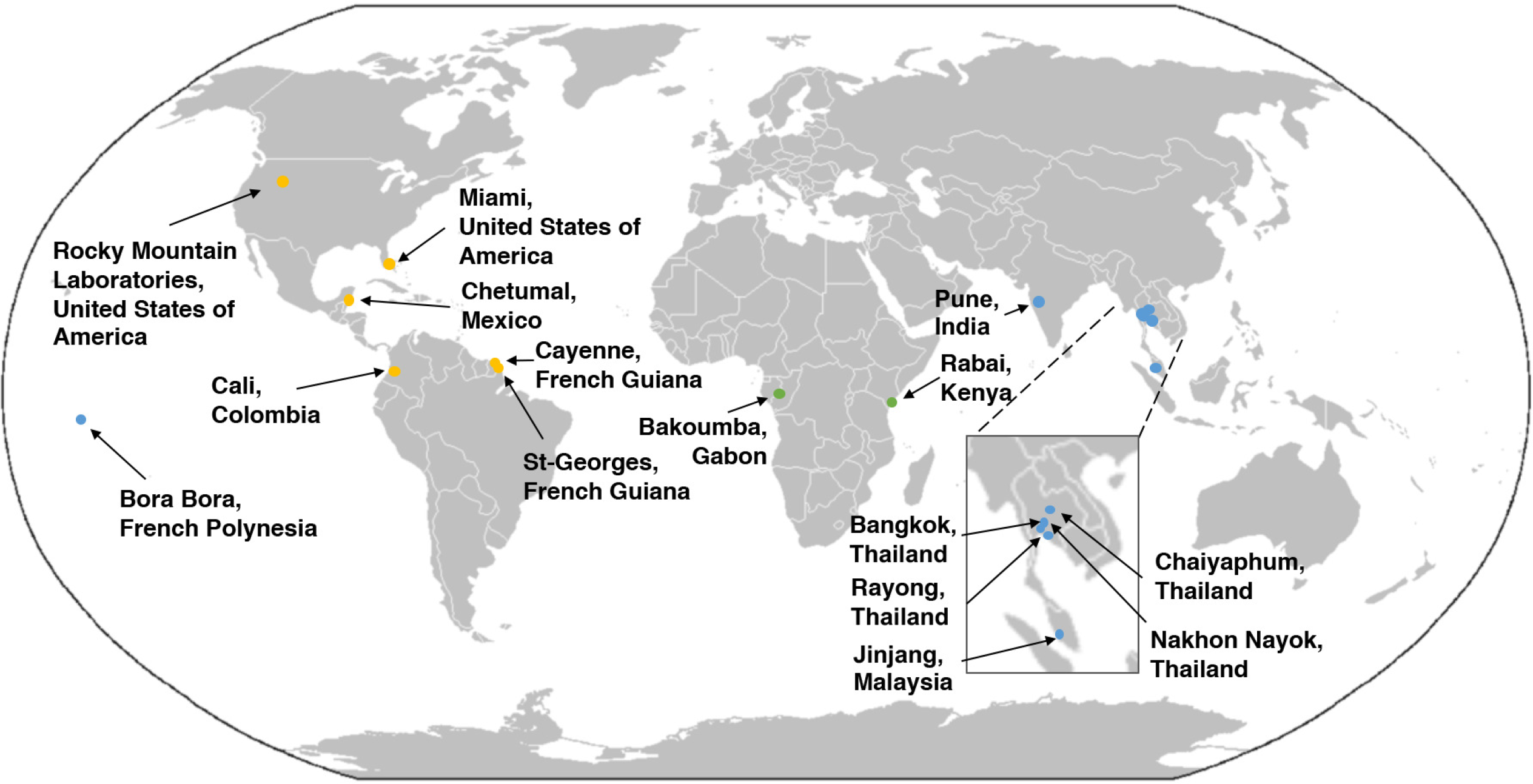
Aedes anphevirus (AeAV) has worldwide distribution in *Ae. aegypti* laboratory colonies, cell lines and wild-caught mosquitoes. Locations of mosquito collection from RNA-Seq data that were positive for AeAV (Table S1). Points refer to collection sites from American (orange), Asia-Pacific (blue) and African (green) locations.

AeAV was present in colonies of *Ae. aegypti* established from eggs collected in Bakoumba, Gabon (48) and also from Rabai, Kenya (designated K2, K14) as well as four mated hybrid strains (49). In colonies wild caught from locations in the Americas (47), full genomes of AeAV were assembled from Miami, USA, Cali, Colombia (40), and Chetumal, Mexico (50, 51) laboratory strains. Partial genomes of AeAV were assembled from Cayenne and St-Georges, French Guiana (52). AeAV was identified in colonies established from eggs collected in Chaiyaphum and Rayong, Thailand (49, 53) as well as Jinjang, Malaysia (54). AeAV was also identified from the widely used Bora-Bora reference strain from French Polynesia (55). AeAV was also present in eight pools of wild-caught *Ae. aegypti* mosquitoes used for ZIKA bio-surveillance in Miami, Florida (56) as well as Nakhon Nayok, and Bangkok, Thailand (18, 22).

In *Aedes* cell lines, AeAV was assembled from RNA-Seq data from the larval *Ae. aegypti* line CCL-125 originally produced in Pune, India (57) and sequenced by the Arthropod Cell Line RNA-Seq initiative, Broad Institute (broadinstitute.org). With the exception of RNA-Seq data from the three *Aedes* cell lines stably infected with *Wolbachia* (RML-12. *w*MelPop-CLA, C6/36. *w*MelPop-CLA and Aag2. *w*MelPopCLA), AeAV was not identified in any other available C6/36 or Aag2 RNA-Seq libraries.

### Genetic variation and evolution of AeAV strains

To assess relatedness and evolution between AeAV strains, a Maximum likelihood phylogeny (PhyML) was undertaken of the CDS region of all strains with complete genomes (Fig. 5A). The unrooted radial phylogenetic tree indicated three strongly supported monophyletic lineages associated with the geographic origin of the sample. We have designated these lineages of AeAV as African, American and Asia-Pacific (Fig. 5B).

**Figure 5.**
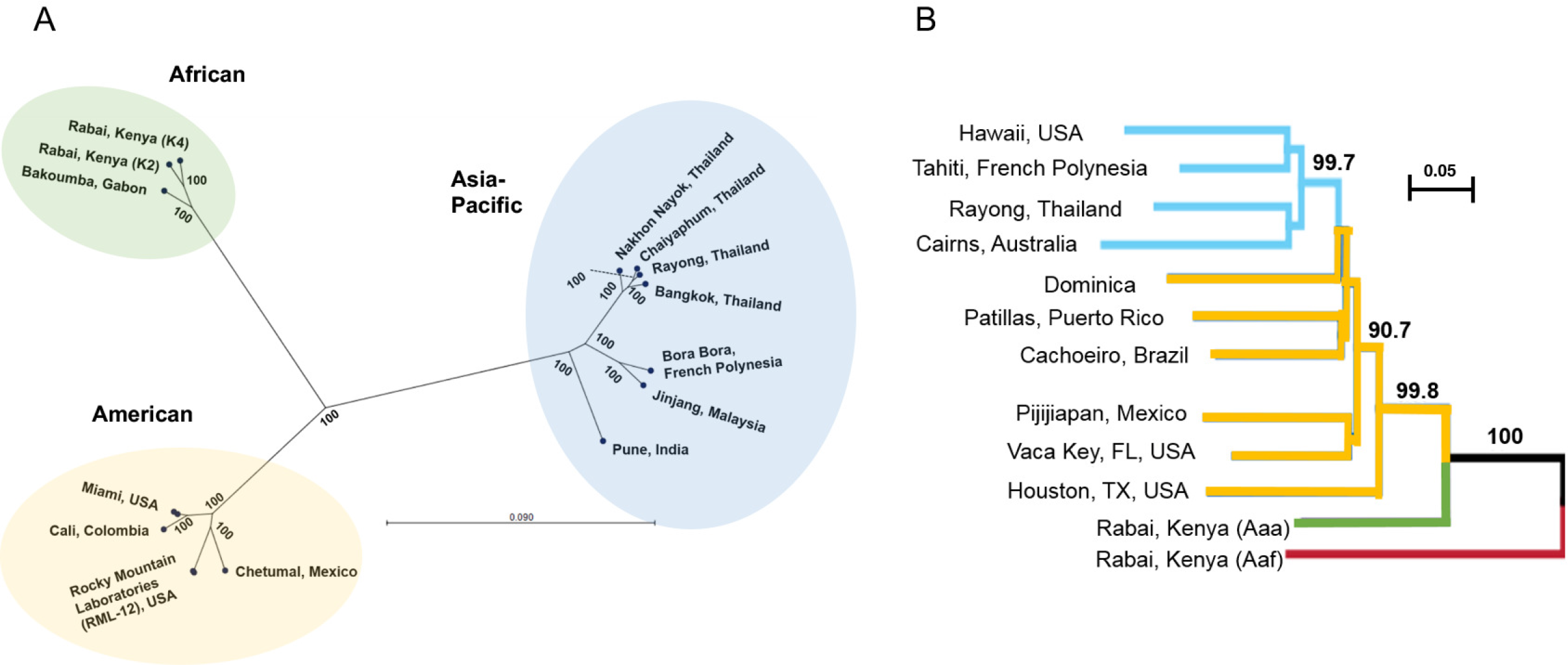
Aedes anphevirus (AeAV) strains have evolved into African, Asia-Pacific and American lineages. **A)** Maximum likelihood phylogeny (PhyML) between AeAV strains using a General Time Reversible (GTR) + G +T model with 1000 bootstraps. Branch lengths represent expected numbers of substitutions per nucleotide site. For visual clarity, the RML-12 clade and Miami clades were collapsed and single examples were shown. **B)** Evolutionary history of worldwide sampling of *Ae. aegypti* adapted from (59, 60) from 1504 SNPs species. Bootstrapped neighbour-joining network based on population pairwise chord-distances from with node support over 90% is shown on relevant branches. New World (American) populations in yellow, and Asia-Pacific populations are shown in light blue. We have truncated the tree and rooted to the *Ae. aegypti formosus* (Aef) shown as a red branch.

In the American lineage of AeAV, all strains that are associated with *Wolbachia*-infected Aedes cell lines (RML-12. *w*MelPop-CLA, C6/36. *w*MelPop-CLA and Aag2. *w*MelPopCLA) are almost identical (99.55-99.86% identity), supporting the hypothesis that contamination of C6/36 and Aag2 cell lines infected with *Wolbachia* is likely from the original RML-12 cell line. AeAV from the eight wild-caught pools of *Ae. aegypti* mosquitoes from Florida, USA (56) and the laboratory colony established from wild collected samples Florida (47) were almost identical (99.86% pairwise identity) with only 17nt differences over the CDS region. The three African lineage strains of AeAV were slightly closer in pairwise nucleotide identity to the American strains (92.65-93.15%) than the Asia/Pacific strains (91.63%-91.74%). All samples that originated from Thailand form a monophyletic group and are closely related to other Thai strains (99.23-99.62%).

We hypothesised that AeAV may have been harboured as part of the virome of *Ae. aegypti* mosquitoes as *Ae. aegypti* expanded from its sub-Saharan African location into the Americas and Asia-Pacific (58). Phylogenetic studies of the *Ae. aegypti* genome support the origin of *Ae. aegypti* from Africa into the New World (Americas) and a subsequent secondary invasion of *Ae. aegypti aegypti* from the New World to the Asia-Pacific region (59, 60). Comparing the evolution of the *Ae. aegypti* nuclear genome with the evolution of AeAV indicates that the Asia-Pacific strains of AeAV have not evolved from the currently circulating American strain lineage. This may indicate that the virus was established independently in both the New-world Americas and also in the Asia-Pacific (Fig. 5B).

### Anphevirus-like insertions into the *Ae. aegypti* genome

The *Ae. aegypti* genome has a large repertoire of virus genes and partial viral genomes, termed Endogenous Viral Elements (EVEs) (61, 62). To explore the possibility of anphevirus-like insertions within the *Ae. aegypti* genome, we queried the most recent Liverpool genome (Aaegl5) with the Aag2. *w*MelPop-CLA AeAV reference strain using the VectorBase BLASTN suite (https://www.vectorbase.org/blast). There were numerous hits of nucleotide similarity (67-70%) of 500-1704nt regions dispersed throughout the *Ae. aegypti* genome. EVEs are acquired through recombination with long terminal repeat (LTR) retrotransposons (62). We present one ~20kb portion of Chromosome 2 of the *Ae. aegypti* genome (Fig. 6) with four anphevirus-like insertions and close proximity to LTR retrotransposable fragments in unidirectional orientation. This suggests insertion of viral elements through LTR retrotransposases and a long evolutionary history of challenge with anphevirus-like species in *Ae. aegypti*.

**Figure 6.**
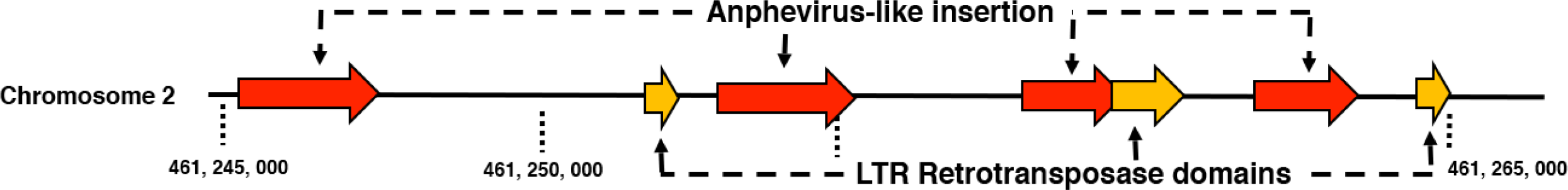
Genomic context for anphevirus-like insertions into the *Ae. aegypti* genome. A 21, 242nt portion of chromosome 2 depicting anphevirus insertions (red) with predicted ORFs that encode for LTR retrotransposase elements (yellow).

### Aedes Anphevirus (AeAV) replicates in *Aedes* cell lines but does not replicate in three mammalian cell lines

Supernatant of Aag2. *w*MelPop-CLA cells was infectious to both *Ae. aegypti* cells (Aa20), and *Ae. albopictus* C6/36 cells over a five-day time course through RT-qPCR analysis (Fig. 7A). Generally, there was significantly more relative AeAV genome copies detected in C6/36 cells at 1 and 5 dpi compared to Aa20 cells. There were also significantly more anti-genome copies of AeAV in C6/36 cells over the five-day time course. The higher replication of AeAV in C6/36 cells as compared to Aa20 cells is not unexpected since C6/36 cells are RNAi deficient and generally RNA viruses replicate more efficiently in the cells (63–65).

**Figure 7.**
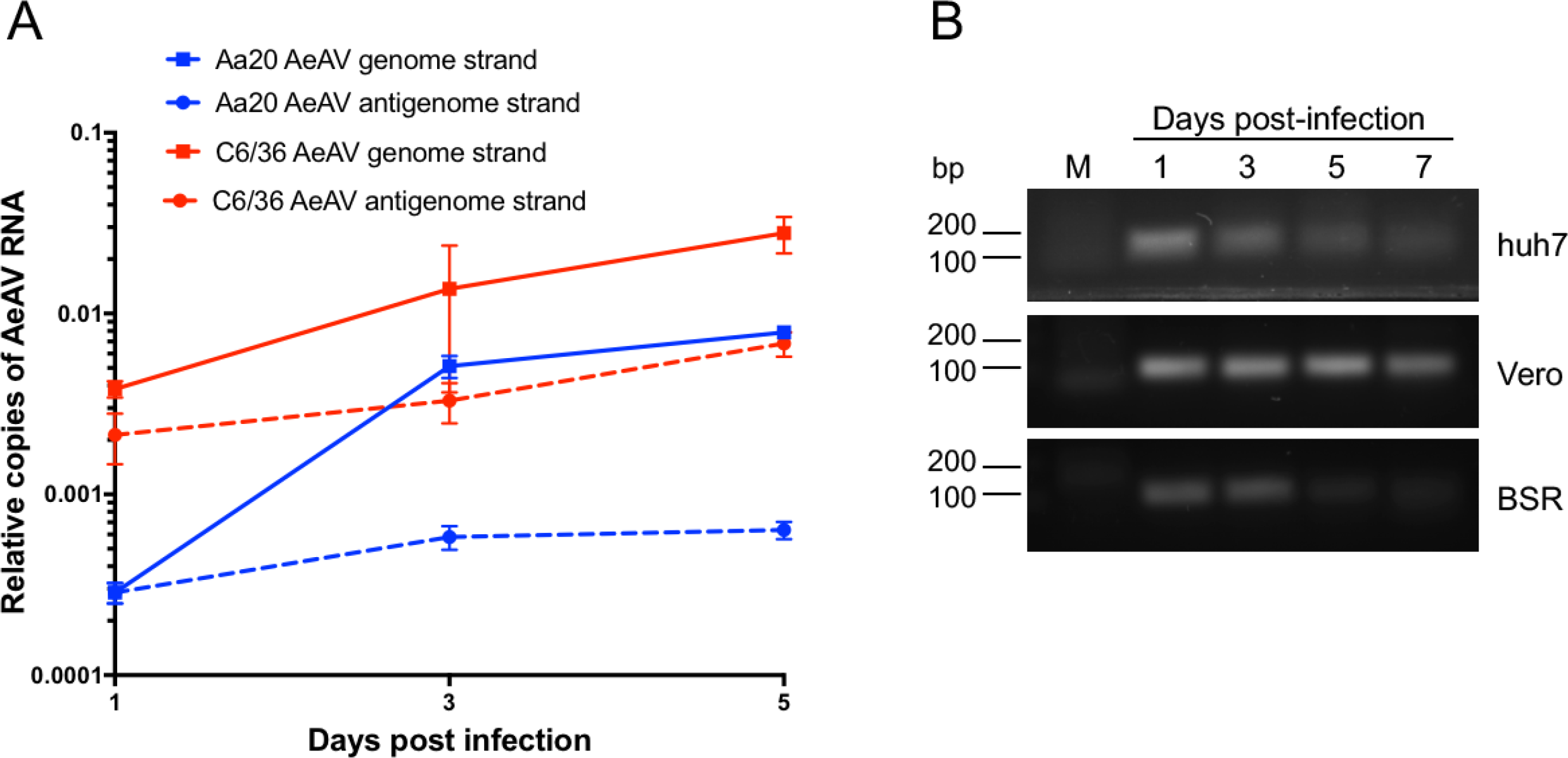
Aedes anphevirus (AeAV) is infectious to *Aedes* cell lines but does not replicate in Huh-7, Vero and BSR vertebrate cell lines. **(A)** RT-qPCR of AeAV genome and anti-genome in a five-day time course in *Ae. aegypti* Aa20 cells, and *Ae. albopictus* C6/36 cells. Error bars represent the SEM of three biological replicates. **(B)** RT-PCR of AeAV genome in a seven-day time course in Human hepatocellular carcinoma cells (Huh-7), African green monkey cells (Vero), Baby Hamster Kidney (BSR). M, Mock infected cells.

We assumed that AeAV is an insect-specific virus based on its phylogenetic position, however, to test if AeAV can replicate in mammalian cells, we inoculated human hepatocellular carcinoma cells (Huh-7), African green monkey cells (Vero), and baby Hamster Kidney (BSR) cells with medium from AeAV-infected cells and performed RT-PCR for AeAV RNA genome abundance over a 7-day time course. While AeAV RNA (most likely from the inoculum) could be detected by RT-PCR at day 1 and 3 after inoculation, it did not increase over time and was visibly reduced in the mammalian cells by 5/7 dpi (Fig. 7B). AeAV was also not detected in the *An. gambiae* cell line MOS-55 transinfected with *w*MelPop-CLA from RML-12-*w*MelPop-CLA (36) sequenced by the Arthropod Cell Line RNA-Seq initiative, Broad Institute (broadinstitute.org). Taken together, the results suggest that AeAV infection may be restricted within the subfamily *Culicinae* or even the *Aedes* genus and is insect specific.

### *Wolbachia pipientis* infection in *Ae. aegypti* cells enhances AeAV replication

As *Wolbachia* is being deployed in the field to reduce dengue transmission, we were interested to find out if it has any effect on replication of AeAV. We extracted RNA from Aag2. *w*MelPop-CLA cells and a previously tetracycline cured Aag2.*w*MelPop-CLA cell line (66), and tested the effect of *Wolbachia* infection on AeAV genome and anti-genome copies. AeAV genomic RNA copies were significantly greater in *Wolbachia*-infected (Aag2.*w*MelPop-CLA) cells than those in tetracycline cleared (Aag2.*w*MelPop-CLA.Tet) *Ae. aegypti* cells, however, there was no statistically significant difference between the relative anti-genome copies between the two cell lines (Fig. 8A).

**Figure 8.**
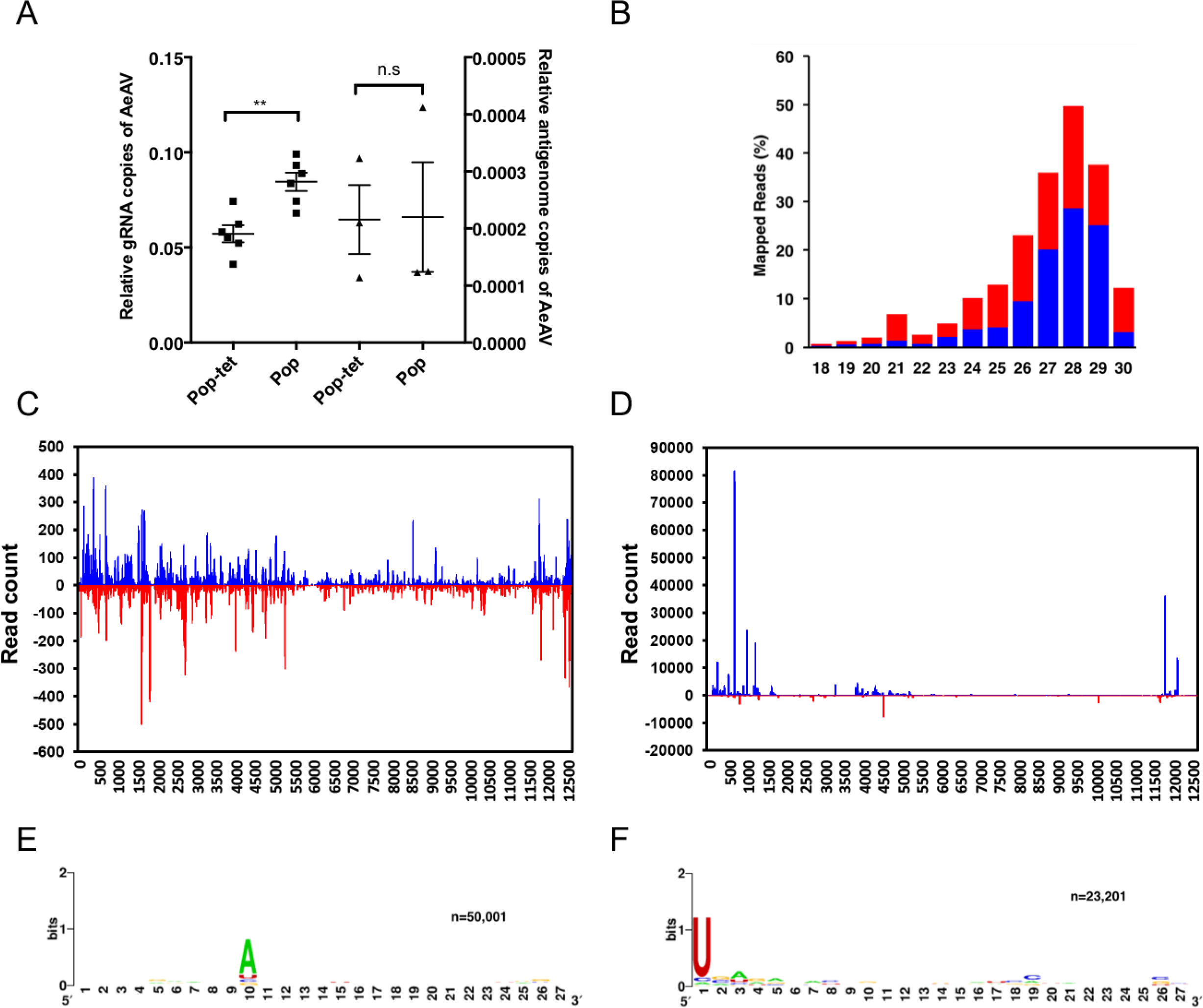
Aedes anphevirus (AeAV) genome replication is enhanced by *Wolbachia* infection in *Ae. aegypti* cells and produces abundant vsiRNAs and vpiRNAs. **A)** RT-qPCR of the AeAV genomic (gRNA) and antigenomic RNA in tetracycline cured Aag2. *w*MelPop-CLA cells (Pop-tet) and Aag2. *w*MelPop-CLA cells (Pop) relative to RPS17. Error bars represent the SEM of six (genome) and three (antigenome) biological replicates. n.s, not significant; **, p < 0.01. **B)** Mapping profile of pooled small RNA fraction in Aag2.*w*MelPop-CLA cells. **C)** Alignment of the 21-nt sRNA reads (representing siRNAs), and D) the 26-31nt reads (representing piRNAs) mapped to the AeAV antigenome (blue) and genome (red) in Aag2.*w*MelPop-CLA cells. Relative nucleotide frequency and conservation of the 28nt small RNA reads that mapped to the **E)** genome, and the **F)** antigenome of AeAV in Aag2.*w*MelPop-CLA cells.

To explore the host small RNA response to AeAV, clean reads from previously prepared sRNA libraries from Aag2. *w*MelPop-CLA and Aag2 (34) were mapped to the AeAV genome. In the cytoplasmic fraction of the Aag2.*w*MelPop-CLA sample 870,012 of 4,686,954 reads (18.56%) mapped to AeAV. In the nuclear fraction 420,215 of 11,406,324 reads (3.68%) mapped to the genome. In the combined Aag2 sRNA library, of 8,600,821 clean reads only four reads mapped to the AeAV genome. The mapping profile of 18-31nt reads mapped from the Aag2. *w*MelPop-CLA library to AeAV indicates a higher proportion 27-31nt viral derived PIWI RNAs (vpiRNAs) than the 21nt vsiRNAs (Fig. 8B).

Analysis of the profile of mapped AeAV vsiRNAs fairly ubiquitously targeted the AeAV genome and anti-genome (Fig. 8C). Hotspots in the vpiRNA mapping profile appeared to target the 3’UTR and ORF1 and the 5’UTR of the AeAV anti-genome (Fig. 8D). Biogenesis of vpiRNAs are independent of Dicer-2 with a bias for adenosine at position 10 in the sense position and a uracil in the first nucleotide of antisense polarity (67). This “ping-pong” characteristic signature was apparent in the vpiRNA reads from the cell line (Fig. 8E and F).

### Persistent infection of AeAV in Aa20 cells modestly reduces replication of DENV-2 genomic RNA

Recently, it has been demonstrated that in *Aedes* cell lines experimentally infected with two ISVs replication of DENV and ZIKV was reduced (13). To test if there was any interaction between AeAV and the subsequent challenge of cells with DENV, we generated an Aa20 cell line inoculated with medium from RML-12 and maintained it for three passages. Aa20 cells persistently infected with AeAV were challenged with 0.1 and 1 multiplicities of infection (MOI) of DENV-2 ET-100 strain. RT-qPCR analysis of DENV-2 genomic RNA showed that accumulation was on average less in AeAV infected Aa20 cells as compared to the control (Fig. 9A and B). This reduction in DENV-2 genome copies was statistically significant at MOI of 0.1 at both three and five days post infection. No AeAV-related RT-qPCR product was detected in mock-infected Aa20 cells (data not shown). We also examined the effect of DENV infection on AeAV RNA levels in the AeAV persistently infected cells. RT-qPCR analysis showed no significant effect on AeAV levels between 1 and 3 days post-DENV infection, however, AeAV genomic RNA levels significantly declined at 5 days-post DENV infection (Fig. 9C). There was no significant difference in the results between 0.1 and 1 MOI of DENV.

**Figure 9.**
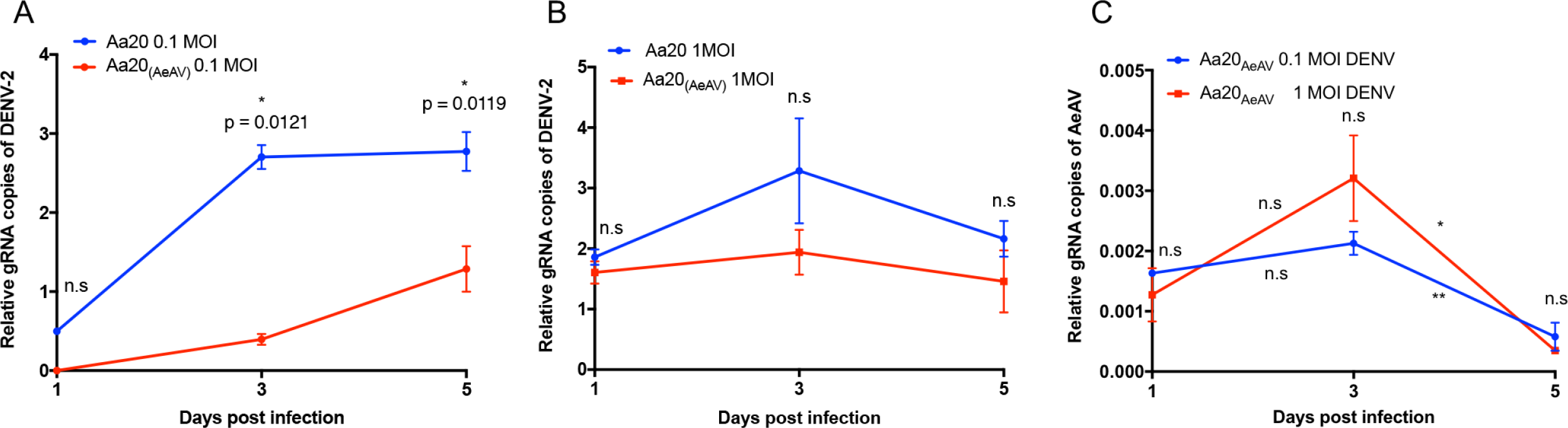
Aedes anphevirus (AeAV) reduces dengue virus replication in Aa20 cells. Aa20 cells persistently infected with AeAV were infected with **(A)** 0.1 and **(B)** 1 MOI of dengue virus serotype 2 (DENV-2). Total RNA was extracted at 0, 1, 3 and 5 days following DENV-2 inoculation and analysed by RT-qPCR. **(C)** RT-qPCR analysis of AeAV persistently infected Aa20 cells infected with 0.1 and 1 MOI of DENV-2 using specific primers to the AeAV genome. Error bars represent the SEM of three biological replicates. n.s, not significant; *, p < 0.05; **, p < 0.01.

### Evidence for vertical transmission of AeAV

We were fortunate to explore the potential vertical transmission of AeAV by using RNA-Seq data of uninfected and infected mated individuals from a study characterising the genetic basis of olfactory preference in *Ae. aegypti* (49). Briefly, McBride and collegues used eggs from a number of *Ae. aegypti* species in Rabai, Kenya to establish laboratory colonies for RNA-Seq analysis. We identified AeAV in the domestic K2 and K14 colonies, which was seemingly absent from the other Rabai (K18, K19, K27) colonies. The K27 colony was interbred with the strain K14, which we found to be AeAV positive. In all the four resultant hybrid colonies, which were subjected to RNA-Seq analysis, we were able to *de novo* assemble identical K14 AeAV strain genomes (Fig. 10).

**Figure 10.**
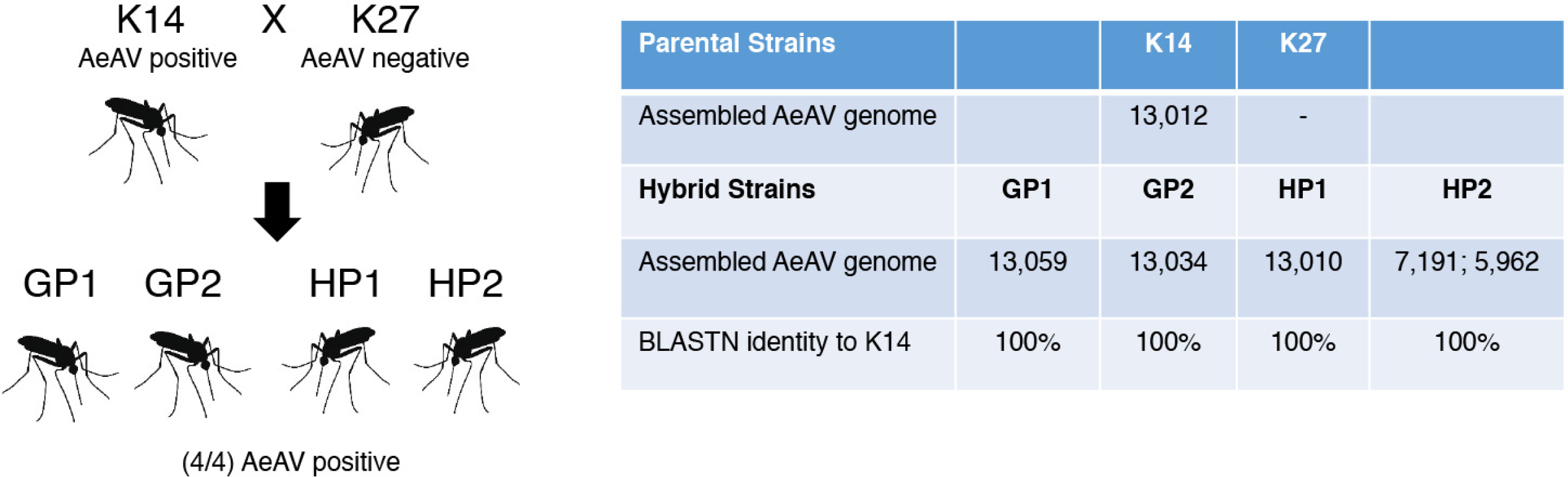
Aedes anphevirus (AeAV) is potentially vertically transmitted. **A)** Diagram showing the parental (K14, K27) and hybrid strains (GP1, GP2, HP1, HP2) from (49). **B)** Table showing assembly statistics and BLASTN similarity of AeAV assembled from K14 and K27 hybrid strains.

The possibility of vertical transmission also is supported by the presence of AeAV in both RNA-Seq data from the sperm of adult male mosquitoes (54) and the female reproductive tract (53).

## Discussion

The ability of AeAV to propagate in *Ae. aegypti* and *Ae. albopictus* cell lines but not in the three mammalian cell lines suggests that AeAV is most likely an ISV, although this needs to be further confirmed using cell lines from other species. To the best of our knowledge, this is the first comprehensive characterisation of any *Anphevirus* species within *Mononegavirales* and the first *Mononegavirales* virus species within *Ae. aegypti*. Although as this manuscript was under revision, the complete genome sequence of AeAV and its phylogenetic relationship with other ISVs was published in a short communication (68). While we have demonstrated that AeAV is spread worldwide in *Ae. aegypti* mosquitoes, we have limited understanding as to how prevalent AeAV is in individual mosquitoes in wild populations, its tissue tropism, or potential impacts on the host. Although it is likely that AeAV is maintained in wild populations of *Ae. aegypti* mosquitoes through vertical transmission, it is possible that AeAV could be maintained though venereal transmission as we were able to identify the whole AeAV genome from a dataset prepared from the sperm of adult male *Ae. aegypti* mosquitoes (54).

To our knowledge, the oldest continually maintained colony of laboratory mosquitoes with AeAV present comes from Jinjang, Malaysia, which was established from wild-collected samples from as early as 1975 (54). This suggests vertical transmission rates of AeAV are high or a high incidence of AeAV within the colony. In comparison, in both French Guiana colonies, we were unable to recover the complete genomes from these strains. It is unlikely that this is due to insertion of AeAV into the nuclear *Ae. aegypti* genome as numerous reads mapped to the ORF7/L region of the reference strain; however, there was not enough coverage to reach consensus of the full genome. As these libraries were prepared from homogenates of mosquitoes, it seems likely that the incidence of infection within these colonies may be lower; however, further analysis would have to be conducted.

In our analysis, AeAV was not detected in any of the widely used Liverpool (LVP) and Rockefeller/UGAL, as well as derived “white eye mutant” strains of *Ae. aegypti*. Analysis of published ncRNA-Seq data from Australian Townsville and Cairns colonies of wild-caught Australian mosquitoes (69, 70) suggests that there is no RNAi response or presence of AeAV in these mosquitoes. Evidence from this study and others suggests that widely used laboratory strains of *Ae. aegypti* harbour a diverse and heterogeneous virome composition and may contribute to the variable vector competence between these colonies.

In our analysis, the geographic origin of the RNA-Seq samples matched the resulting phylogenetic relationship of each strain. The presence of AeAV in the *Ae. albopictus* cell line RML-12, presumably the origin of the AeAV contamination in other *Aedes* cell lines transinfected with adapted *w*MelPop strain, was the only *Ae. albopictus* sample in our analysis. During our analysis, we queried all of the 266 currently available *Ae. albopictus* RNA-Seq datasets (TaxonID: 7160) uploaded to the SRA, none of which indicated presence of AeAV. We hypothesise that AeAV from RML-12 is likely due to contamination as the cell line is often mischaracterised as originating from *Ae. aegypti* (71, 72). In many laboratories that study arbovirus interactions, more than one *Aedes* cell line is maintained. As the RML-12 AeAV strain is genetically placed within the American lineage, and the namesake of the cell line, Rocky Mountain Laboratories, (71) located in Montana, USA suggests possible contamination from domestic *Ae. aegypti* mosquito samples. While no other *Ae. albopictus* RNA-Seq data were positive for AeAV in this study, we cannot rule out the possibility that AeAV could exist in American populations of *Ae. albopictus. Ae. aegypti* and *Ae. albopictus* co-exist in North America and compete for larval habitats of discarded tires and other artificial containers (73).

RNAi response is commonly observed in mosquitoes against RNA viruses. This response includes miRNA, siRNA and piRNA pathways (74, 75). Similarly, we found a large number of 21nt vsiRNAs produced against AeAV in infected cells that were evenly mapped to both sense and antisense strands, indicating that dsRNA intermediates produced during replication must be the target of the host cell RNAi response. In addition, a large number of vpiRNAs were found mapped to the 5’UTR, ORF1 and the 3’UTR of the AeAV genome. These vpiRNAs had the typical ping-pong signature (U_1_-A_10_) of secondary piRNAs. This signature has also been found in vpiRNAs produced during alphavirus (76) and bunyavirus (63, 77) infections, but not in vpiRNA-like small RNAs in most flaviviruses, such as DENV (78), ZIKV (79, 80) and an insect-specific flavivirus (81). We found a higher proportion of small RNAs from Aag2-*w*MelPop-CLA cells that mapped to AeAV are vpiRNAs (about 50%) as compared to less than 10% vsiRNAs. Literature suggests that when the siRNA pathway is compromised more vpiRNAs are produced. This has been shown in RNAi deficient C6/36 cells when infected with Sindbis virus, Rift Valley fever virus or La Crosse virus (63–65). The RNA-Seq data from C6/36. *w*MelPop-CLA cells were for long transcripts rather than small RNAs, therefore we were not able to confirm if in those cells there are higher proportion of vpiRNAs than vsiRNAs. The over-representation of vpiRNAs in respect to vsiRNAs has also been demonstrated in negative-sense *Bunyavirales* members PCLV and RVFV (30, 65) in Aag2 cells for all segments of the genome. It remains to be seen however if the higher vpiRNA to vsiRNA ratio in Aag2. *w*MelPop-CLA could be due to suppression of the siRNA pathway by AeAV, or alternatively, *Wolbachia* may have an effect on the siRNA pathway. It seems however more likely that as negative sense RNA viruses produce less dsRNA replicative intermediates, these could be simply less targeted by the siRNA pathway and are unable to be resolved by sRNA profiling. These possibilities require further investigations, and it remains to determine the role of the vpiRNAs in AeAV replication or host anti-viral response.

The effects of *Wolbachia* on virus restriction are variable and depend on *Wolbachia* strain, virus family and transinfected host (82). *Wolbachia* was shown to enhance AeAV replication in *Ae. aegypti* cells in this study. Recent studies have demonstrated that *Wolbachia* has no effect on multisegmented negative-sense RNA viruses; for example, Phasi Charoen-like virus (PCLV) (Family: *Bunyaviridae*) in *w*MelPop-CLA strain infected Aag2 cells (30), La Crosse virus (Family: *Bunyaviridae*) and vesicular stomatitis virus (Family: *Rhabdoviridae*), which is in the same order as AeAV, in *w*Stri strain in *Ae. albopictus* C710 cell line (83), and the Rift Valley fever virus (RVFV) (Family: *Phenuiviridae*), in *Culex tarsalis* mosquitoes transinfected with a somatic *Wolbachia* (strain wAlbB) had no effect on RVFV infection or dissemination (84), respectively. RVFV and PCLV belong to the *Bunyavirales* order and AeAV belongs to *Mononegavirales*, distinct orders of negative strand RNA viruses. All three viruses have conserved features that may provide insight into how they might be protected from restriction by *Wolbachia*. The genome of all negative-sense ssRNA viruses is both encapsidated within the nucleoprotein (85, 86) and is attached to RNA dependent RNA polymerase within the virion (87). The RNA dependent polymerase complex carries out both transcription of virus genes and replication of the genome.

*Wolbachia* has been shown to restrict a number of positive-sense RNA virus species from the *Togaviridae* and *Flaviviridae* families (82). After fusion and entry into the host cell, the genomes of *Togaviridae* and *Flaviviridae* species are released into the cytoplasm and translated directly into polypeptide protein(s). These polypeptide proteins are processed by viral and cellular proteases to generate the mature structural and non-structural proteins which are then used to replicate the genome (88). While the exact mechanism for RNA virus restriction in *Wolbachia*-infected insects has remained elusive, it has been shown that restriction of RNA viruses by *Wolbachia* happens early in infection (89, 90). In the *Ae. albopictus* cell line C710 stably infected with wStri, the polypeptide of ZIKV is not produced as determined by immunoblot at one day post-infection (90). Additionally, *Wolbachia* exploits host innate immunity to establish a symbiotic relationship with *Ae. aegypti* (91). Perhaps the combination of protection of the RNA nucleocapsid genome or genome segments when released in the cytoplasm or activity of the RNA-dependent RNA polymerase may aid in evasion of host immune response enhanced by *Wolbachia* or *Wolbachia* effector molecules (92). However, a recent study suggested increases in infection of *Ae. aegypti* mosquitoes by insect-specific flaviviruses when they harbour *Wolbachia* wMel strain (93).

The ability of AeAV to modestly reduce DENV-2 genomic RNA in a persistently infected cell line was unexpected. While it has previously been shown that members of the same virus family can provide super exclusion of additional viruses (12, 94), little work has been undertaken to look at cross viral family exclusion effects. Our results showed that less DENV replication occured in the presence of AeAV, with the difference particularly significant at lower MOI. If this suppressive effect also occurs in mosquitoes, enhancement of AeAV in *Ae. aegypti* mosquitoes infected with *Wolbachia* may be beneficial in terms of DENV suppression.

As *Ae. aegypti* is perhaps the most important vector of arboviruses worldwide, further work should be undertaken in understanding and characterising the virome of this mosquito and effects on mosquito life-history traits. Our findings provide new insights into the evolution and genetic diversity of AeAV across a wide geographic range as well as providing valuable insights into the virus features and families restricted by *Wolbachia* in mosquito hosts and its effects on arboviruses they transmit.

## Materials and Methods

### Cell lines maintenance and experimental infection with AeAV

*Ae. aegypti* cell line (Aag2) stably infected with *Wolbachia* (denoted Aag2.*w*MelPop-CLA) as previously described for the C6/36.*w*MelPop-CLA (35), with its previously generated tetracycline-treated line (66), and both *Ae. albopictus* C6/36 cell line (57) and RML-12 cell lines were maintained in 1:1 Mitsuhashi-Maramorosch and Schneider’s insect medium (Invitrogen) supplemented with 10% Fetal Bovine Serum (FBS, Bovogen Biologicals). Aa20 cells established from *Ae. aegypti* larvae (95) were maintained in Leibovitz’s L15 medium supplemented with 10% FBS (France - Biowest) and 10% Tryptose phosphate broth at 27°C. African green monkey cells (Vero) were maintained in Opti-MEM I Reduced-Serum Medium supplemented with 2% FBS and 10 mL/L Penicillin-Streptomycin (Sigma-Aldrich). Human hepatocellular carcinoma cells (Huh-7) were maintained in Opti-MEM I Reduced-Serum Medium supplemented with 5% FBS and 10 mL/L Penicillin-Streptomycin (Sigma-Aldrich). BSR cells (a clone of Baby Hamster Kidney-21) were maintained in Dulbecco’s Modified Eagle Medium (Gibco), 2% FBS and 10 mL/L Penicillin-Streptomycin (Sigma-Aldrich). All mammalian cells were kept at 37°C with 5%CO_2_.

For experimental infection of cells, 10^6^ cells of *Aedes* or mammalian cells were seeded in a 12-well plate. Subsequently, supernatant from Aag2.*w*MelPop-CLA cells was collected, centrifuged at 2150xg for 5 min to remove cells and debris, and used as an AeAV inoculation source. One Aa20 cell line was experimentally inoculated with RML-12 cell supernatant and kept as a persistently infected AeAV cell line. Cells were collected at 1, 3 and 5 days post-inoculation for *Aedes* cell lines for RT-qPCR analysis and 1, 3, 5 and 7 days for mammalian cell lines for RT-PCR analysis.

### Aedes anphevirus (AeAV) and dengue virus (DENV-2) interaction assay

The third passage of Aa20 cells persistently infected with AeAV (denoted Aa20AeAV) and Aa20 mock were seeded at the density of 3×10^5^ cells in 12-well plates overnight. Cells were then infected with the East Timor (ET-100) DENV-2 strain at a multiplicity of infection (MOI) of 0.1 and 1, cells were rocked for an hour at room temperature and supernatant was discarded and replaced with fresh medium. Cells were collected at 0, 1, 3 and 5dpi after infection with mock collected at 5 dpi. Cells were subjected to RNA extraction to quantify the DENV-2 genomic RNA levels by RT-qPCR as described below.

### Assembly and identification of AeAV strains from RNA-Seq data

For detection of AeAV in previously published RNA-Seq data, we used the assembled RML-12 AeAV genome as a BLASTN query for all available *Ae. aegypti* (taxonID: 7159) RNA-Seq data within the Sequence Read Archive (SRA) on NCBI. SRA run files with positive hits of 90-100% identity and an E value <2E-30 were downloaded and converted to fastq using the NCBI SRA toolkit https://www.ncbi.nlm.nih.gov/sra/docs/toolkitsoft/ for further analysis. FastQC (https://www.bioinformatics.babraham.ac.uk/projects/fastqc/) was used for quality checking of fastq files and adapter identification. Fastq files were then imported into CLC Genomics Workbench (10.1.1) and were adapter and quality trimmed (<0.02; equivalent Phred quality score of 17, ambiguous nucleotides: 2).

Two strategies were used to assemble strains of AeAV; fastq files from the same source of *Ae. aegypti* were pooled and *de novo* assembled using the CLC Genomics Workbench assembly program with automatic bubble and word sizes. This was sufficient to assemble the full coding sequences (CDS) of most strains of AeAV. Table S1 contains *de novo* assembly statistics from each dataset used.

If *de novo* assembly did not produce the complete AeAV genome, to complete further sections of the AeAV genome clean reads were mapped to the C6/36.*w*MelPop-CLA strain of AeAV with stringent alignment criteria (Match score:1 Mismatch cost:2 length fraction 0.89 and similarity fraction 0.89) to exclude false-positive mapping that derives from Endogenous Viral Elements (EVEs). To confirm accuracy of assembly, the largest contigs of consensus mapping were extracted and then used as a reference for re-mapping and manually checked. Final sequences of the virus genomes were obtained through the majority consensus of the mapping assembly and were given Coding Complete (CC) or Standard Draft (SD) genome quality ratings (96).

### RNA isolation, strand specific cDNA synthesis and RT-qPCR

Total RNA was extracted from mosquito cells using QIAzol Lysis Reagent (QIAGEN) and treated with Turbo DNase (Thermo Fisher Scientific) as per manufacturer’s instructions. RNA quality and quantity were evaluated using a BioTek Epoch Microspot Plate Spectrophotometer. For the production of AeAV genome and anti-genome cDNA, two cDNA reactions were generated using 600ng of RNA and SuperScript III Reverse Transcriptase (Thermo Fisher Scientific). The genome cDNA strand was synthesised using a forward primer to AeAV (AeAVGenome-RT 5’- AGACTTCTAAGCCTGCCCACA -3’), and the AeAV anti-genome cDNA strand was synthesised using a reverse orientation primer (AeAVAntiGenome-RT 5’- ACACTTGCCATGTGCTCAG -3’). *Aedes* Ribosomal protein subunit 17 (RPS17) primers (*Ae. aegypti:* AeRPS17-qR 5’- GGACACTTCGGGCACGTAGT-3’ and *Ae. albopictus* AalRPS17q-R 5’- ACGTAGTTGTCTCTCTGCGCTC -3’) were used for reference gene cDNA synthesis. Following RT, qPCR with AeAV primers (AeAV-qF 5’-GACAATCGCATTGGCTGCAT-3’ and AeAV-qR 5’- CCCGAGACAATCGGCTTCTT -3’) as well as primer pairs for the RPS17 genes (*Ae. aegypti:* AeRPS17-qF 5’- CACTCCGAGGTCCGTGGTAT -3’ and *Ae. albopictus:* AalRPS17-qF 5’-CGCTGGTTTCGTGACACATC -3’) were undertaken. RPS17 was used for normalizing data as described previously in *Ae. aegypti* cells (97).

For quantitation of DENV-2 genome copies in Aa20 cells, two SuperScript III reverse transcription (Thermo Fisher Scientific) reactions with 1000ng of RNA were prepared. For DENV-2 genome copies reaction, the reverse primer (DENV2-qR 5’- CAAGGCTAACGCATCAGTCA -3’) and in a separate cDNA synthesis reaction the *Ae. aegypti* RPS17 primers as described above. Subsequently, qPCR for DENV-2 was carried out using the DENV-2 primer pair (DENV2-qF 5’- GGTATGGTGGGCGCTACTA -3’ and DENV2-qR), and RPS17 was used as a normalising control as also described above.

Each time-point for experimental infection was run with three biological replicates and two technical replicates. Platinum SYBR Green Mix (Invitrogen) was used for qPCR with 20ng of RT products in a Rotor-Gene thermal cycler (QIAGEN) as described above. The relative abundance of AeAV RNA and DENV-2 to the host reference gene was determined by qGENE software using the ΔΔC_t_ method and analyzed using GraphPad Prism.

To test for the replication of AeAV in mammalian cells, 1000ng of total RNA extracted from the cells was extracted and used for first stand synthesis with SuperScript III reverse transcriptase with the AeAVGenome-RT primer. qPCR was subsequently carried out using AeAVGenome-RT primer and the qPCR primer AeAVqR2 5’-ATGAAAGTATGGATACACACCGCG-3’. Products were then visualised on a 1% agarose gel.

### Virus genome annotation

Potential ORFs of AeAV were analysed using NCBI Open Reading Frame Finder (https://www.ncbi.nlm.nih.gov/orffinder/) with a minimal ORF length of 150. ORFs were cross referenced with mapping from Poly(A) enriched transcriptomes (Fig. S1) to reduce false positive identification of ORFs. For determination of putative domains in AeAV, ORFs were translated and searched against the Conserved Domain Search Service (CD Search) (https://www.ncbi.nlm.nih.gov/Structure/cdd/wrpsb.cgi). For protein homology detection, we used the HHPred webserver on translated AeAV ORFs (98).

To best discriminate N-terminus transmembrane domains from signal peptides, we used the consensus TOPCONS webserver (99). Glycosylation sites were predicted by the NetNGlyc 1.0/ NetOGlyc 4.0 Server (http://www.cbs.dtu.dk/services/).

### Phylogenetic analysis

For placement of AeAV within the order *Mononegavirales*, ClustalW was used on CLC Genomics Workbench to align amino acid sequences of 30 L proteins of the most closely related *Mononegavirales* species as determined by BLASTp of the NCBI non-redundant database. A Maximum likelihood phylogeny (PhyML) was constructed using the Whelan and Goldman (WAG) amino acid substitution model with 1000 bootstraps. Accession numbers and associated hosts used to produce the phylogenetic tree are available in Table S1.

To determine relatedness between different strains of AeAV, genomes that were coding complete and greater than 30× coverage were trimmed of 3’UTR and 5’UTR regions and aligned using the ClustalW algorithm on CLC Genomics Workbench. A Maximum likelihood phylogeny (PhyML) was constructed. A Hierarchical likelihood ratio test (hLRT) with a confidence level of 0.01 suggested that the General Time Reversible (GTR) +G (Rate variation 4 categories) and +T (topology variation) nucleotide substitution model was the most appropriate. 1000 bootstrap replicates were performed with 95% bootstrap branching support cut-off.

### Statistical analysis

Statistical analysis Unpaired t-test was used to compare differences between two individual groups, while One-way ANOVA with Tukey’s post-hoc test was carried out to compare differences between more than two groups. Data that did not pass the normality test were re-analysed by the non-parametric Wilcoxon test indicated in their relevant figure legends.

### Accession numbers

All the complete and incomplete virus genome sequences generated in this study have been deposited in Genbank under the accession numbers (MH037149 and MH430648-MH430666).

## Acknowledgements

This project was funded by the Australian Research Council grant (ARC, DP150101782) to SA and University of Queensland scholarship to RP. For the supply of the DENV-2 (ET-100) strain, we thank Dr Daniel Watterson and Professor Paul Young from the University of Queensland. We thank Dr Daniel Watterson and Dr Jody Hobson-Peters for providing Huh-7 and BSR cells, respectively. The authors would like to thank the technical assistance of Dr Sultan Asad, Dr Kayvan Etebari and Hugo Perdomo Contreras, as well as current and former members of the Asgari lab for their fruitful conversations.

